# Attractor Landscape Analysis Distinguishes Aging Markers from Rejuvenation Targets in Human Keratinocytes

**DOI:** 10.64898/2026.02.01.703159

**Authors:** Neil Copes, Clare-Anne Edwards Canfield

## Abstract

Cellular aging is characterized by progressive changes in gene expression that contribute to tissue dysfunction; however, identifying genes that regulate the aging process, rather than merely serve as biomarkers, remains a significant challenge. Here we present PRISM (Pseudotime Reversion via In Silico Modeling), a computational pipeline that integrates pseudotime trajectory analysis with Boolean network analysis to identify cellular rejuvenation targets from single-cell RNA sequencing data. We applied PRISM to a published dataset of human skin comprising 47,060 cells from nine donors aged 18 to 76 years. Analysis of keratinocytes revealed two distinct aging trajectories with fundamentally different regulatory architectures. One trajectory (labeled Y_272) exhibited “aging as convergence,” where cells were driven toward a single dominant aged attractor (aging score +2.181). A second trajectory (labeled Y_308) exhibited “aging as departure,” where cells escaped from a dominant youthful attractor basin (aging score −0.536). Systematic perturbation analysis revealed a critical distinction between genes exhibiting age-related expression changes (phenotypic markers) and genes controlling attractor landscape architecture (regulatory controllers). Switch genes marking the aging trajectories proved largely ineffective as intervention targets, while master regulators operating at higher levels of the regulatory hierarchy produced substantial rejuvenation effects. BACH2 knockdown was identified as the dominant intervention for Y_272, shifting the aging score by Δ= −3.746 (98.9% improvement). ASCL2 knockdown was identified as the top target for Y_308, with synergistic enhancement observed through combinatorial perturbation with ATF6. These findings demonstrate that attractor-based analysis identifies different and potentially superior therapeutic targets compared to expression-based approaches and provide specific hypotheses for experimental validation of cellular rejuvenation strategies in human skin.

## Introduction

Cellular aging represents one of the most fundamental challenges in biology and medicine. As organisms age, their cells undergo progressive changes in gene expression, metabolic function, and regenerative capacity that ultimately contribute to tissue dysfunction and age-related disease (Lopez-Otin et al., 2013). Understanding the molecular mechanisms that drive these changes, and identifying strategies to reverse them, has become a central goal of modern biomedical research. The advent of single-cell RNA sequencing (scRNA-seq) has transformed our ability to characterize aging at cellular resolution. By profiling gene expression in thousands of individual cells, researchers can now identify cell type-specific aging signatures, map cellular heterogeneity within aging tissues, and trace developmental and differentiation trajectories that may be altered with age (Tabula Muris Consortium, 2020). Studies applying these approaches to human tissues have revealed tissue-specific patterns of cellular aging, identified senescent cell populations, and characterized the senescence-associated secretory phenotype (SASP) that contributes to chronic inflammation in aged tissues (Zou et al., 2021; Kimmel et al., 2019). Despite these advances, a fundamental limitation of expression-based approaches remains. These approaches excel at describing what changes during aging but provide limited insight into what controls those changes. Differential gene expression analysis identifies genes whose expression levels differ between young and old cells, but these genes may be downstream effectors of the aging process rather than upstream regulators. The distinction matters profoundly for therapeutic development, as targeting downstream markers is unlikely to reverse the underlying aging program, while targeting upstream controllers could potentially reprogram cells toward more youthful states. Boolean network modeling offers a complementary framework for understanding cellular regulation that addresses this limitation. Originally developed by Kauffman (1969) to model gene regulatory networks, Boolean networks represent genes as binary variables (ON or OFF) connected by logical rules that determine how each gene’s state depends on its regulators. Despite their apparent simplicity, Boolean networks capture essential features of cellular regulation and exhibit rich dynamical behavior, including the emergence of stable configurations called attractors that correspond to distinct cell states or phenotypes (Kauffman, 1993; Huang et al., 2005). The attractor concept provides a powerful lens for understanding cellular identity and state transitions. In the state space defined by all possible combinations of gene expression patterns, attractors represent stable configurations toward which the regulatory network naturally evolves from nearby initial conditions. The basin of attraction surrounding each attractor encompasses all initial states that eventually converge to that attractor. Different cell types, differentiation states, and disease conditions can be understood as distinct attractors in the regulatory landscape, with transitions between states corresponding to movement between attractor basins (Huang et al., 2009; Li et al., 2014). This framework suggests a novel approach to understanding and reversing cellular aging. If aged and youthful cellular phenotypes correspond to distinct attractors in the gene regulatory landscape, then aging can be conceptualized as transitions between attractor basins rather than simple accumulation of molecular damage. More importantly, identifying the regulatory factors that control attractor stability and basin boundaries could reveal intervention targets capable of redirecting cells from aged to youthful attractors, effectively reversing the aging phenotype at the regulatory level.

Here we present PRISM (Pseudotime Reversion via In Silico Modeling), a computational pipeline that integrates pseudotime trajectory analysis with Boolean network modeling to identify cellular rejuvenation targets. PRISM constructs executable Boolean networks from scRNA-seq data by combining trajectory inference, switch gene identification, and transcription factor network analysis, then systematically tests gene perturbations to identify interventions that shift the attractor landscape toward more youthful configurations. We applied PRISM to a published scRNA-seq dataset comprising skin biopsies from nine human donors aged 18 to 76 years (Zou et al., 2021). Our analysis focused on keratinocytes, the dominant cell type in skin, and identified two distinct aging trajectories with fundamentally different attractor architectures. Critically, we found that genes exhibiting the clearest age-related expression changes (phenotypic markers) were largely distinct from genes controlling attractor landscape architecture (regulatory controllers), with the latter representing far more promising therapeutic targets. For each trajectory, we identified master regulators whose perturbation was predicted to produce substantial shifts toward youthful attractor configurations, providing concrete hypotheses for experimental validation.

The objectives of this study were threefold: (1) to demonstrate that Boolean network modeling can capture meaningful aging-related dynamics from scRNA-seq data, (2) to test whether attractor-based analysis identifies different therapeutic targets than expression-based approaches, and (3) to generate specific, experimentally testable predictions for cellular rejuvenation interventions in human skin.

## Results

### Dataset Overview and Cell Type Selection

Single-cell RNA sequencing data were obtained from the dataset published by Zou et al. (2021) through the Human Cell Atlas Data Explorer, comprising skin biopsies from nine human donors aged 18 to 76 years. Quality control and normalization retained 47,060 cells across 25 distinct cell types, identified through SingleR annotation (Aran et al., 2019) using the Human Primary Cell Atlas reference. Keratinocytes dominated the cellular population (Table 1), representing 80.4% of all cells, followed by epithelial cells (5.9%), endothelial cells (2.2%), and fibroblasts (2.1%). Random downsampling ensured equal representation of each donor within the keratinocyte population to prevent age-related batch effects from confounding downstream pseudotime and attractor analyses.

**Table 1:**
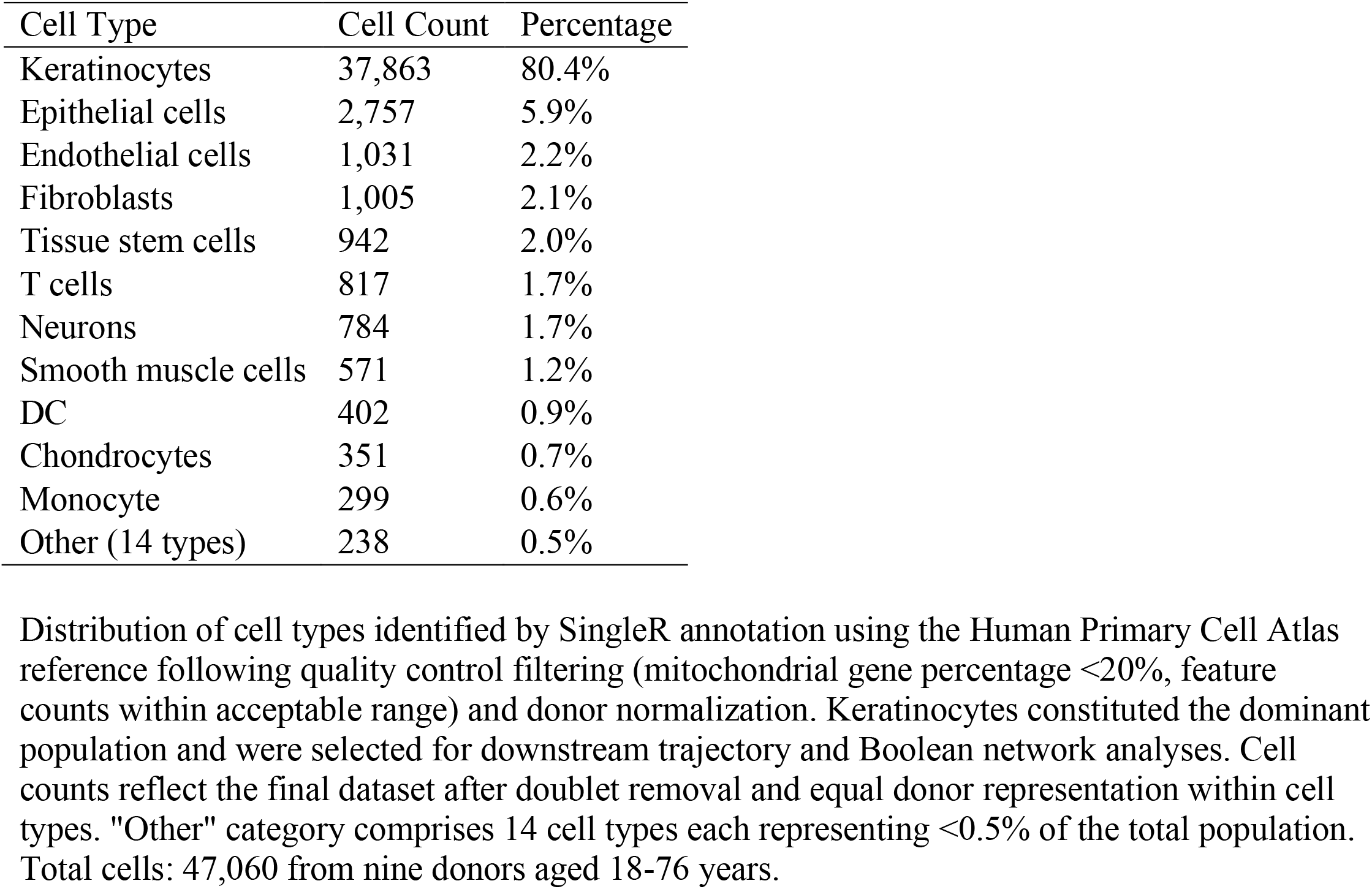
Cell Type Abundance Following Quality Control and Donor Normalization.

| Cell Type | Cell Count | Percentage |
| --- | --- | --- |
| Keratinocytes | 37,863 | 80.4% |
| Epithelial cells | 2,757 | 5.9% |
| Endothelial cells | 1,031 | 2.2% |
| Fibroblasts | 1,005 | 2.1% |
| Tissue stem cells | 942 | 2.0% |
| T cells | 817 | 1.7% |
| Neurons | 784 | 1.7% |
| Smooth muscle cells | 571 | 1.2% |
| DC | 402 | 0.9% |
| Chondrocytes | 351 | 0.7% |
| Monocyte | 299 | 0.6% |
| Other (14 types) | 238 | 0.5% |

Distribution of cell types identified by SingleR annotation using the Human Primary Cell Atlas reference following quality control filtering (mitochondrial gene percentage <20%, feature counts within acceptable range) and donor normalization. Keratinocytes constituted the dominant population and were selected for downstream trajectory and Boolean network analyses. Cell counts reflect the final dataset after doublet removal and equal donor representation within cell types. "Other" category comprises 14 cell types each representing <0.5% of the total population. Total cells: 47,060 from nine donors aged 18-76 years.

### Pseudotime Trajectory Analysis

Monocle3 trajectory inference (Trapnell et al., 2014; Cao et al., 2019) on the 37,863 keratinocytes identified nine distinct pseudotime branches, each representing a potential developmental or differentiation pathway within the keratinocyte lineage. To validate which trajectories represented genuine aging processes rather than other biological programs, Spearman correlation analysis was performed between pseudotime values and donor chronological age for each branch. A conservative threshold (*r* ≥ 0.3, *p* < 0.05) was established, representing the minimum signal-to-noise ratio where aging effects dominate over non-aging factors in the trajectory. Two trajectories exceeded this threshold with highly significant correlations (Table 2 and Figure 1). The trajectory labeled Y_308 demonstrated the strongest age association (*r* = 0.440, *p* = 4.3 × 10^−51^, *n* = 1,053 cells), while the trajectory labeled Y_272 showed robust but slightly weaker correlation (*r* = 0.367, *p* = 9.9 × 10^−41^, *n* = 1,239 cells). Both trajectories maintained representation across all nine donors, ensuring that observed aging signatures reflected genuine biological changes rather than donor-specific artifacts. The remaining seven trajectories showed substantially weaker or non-significant correlations with age (*r* values ranging from 0.046 to 0.246), indicating they represent biological processes orthogonal to aging, such as spatial differentiation programs or cell cycle dynamics. The two validated trajectories exhibited similar temporal characteristics with an expansion of cells at higher pseudotime values among older donors. This expansion was especially pronounced in the oldest donor (76 years old).

**Table 2:** Spearman Correlation Between Pseudotime and Donor Age Across all Keratinocyte Trajectories.

| Trajectory | Correlation (*r*) | *p*-value | Cell Count | Meets Threshold |
| --- | --- | --- | --- | --- |
| Y_308 | 0.440 | $4.3 \times 10^{-51}$ | 1,053 | Yes |
| Y_272 | 0.367 | $9.9 \times 10^{-41}$ | 1,239 | Yes |
| Y_176 | 0.246 | $1.7 \times 10^{-38}$ | 2,700 | No |
| Y_217 | 0.239 | $1.9 \times 10^{-17}$ | 1,233 | No |
| Y_267 | 0.181 | $4.0 \times 10^{-21}$ | 2,669 | No |
| Y_193 | 0.097 | $2.3 \times 10^{-4}$ | 1,432 | No |
| Y_225 | 0.091 | $9.4 \times 10^{-6}$ | 2,352 | No |
| Y_187 | 0.047 | 0.133 | 1,015 | No |
| Y_243 | 0.039 | 0.034 | 3,007 | No |

Correlation analysis validating which of the nine Monocle3-identified trajectories represent genuine aging processes. Trajectories were ranked by Spearman correlation coefficient (*r*) between pseudotime values and donor chronological age. The threshold for inclusion in downstream Boolean network analysis required both statistical significance ( $p < 0.05$ ) and minimum effect size ( $r \geq 0.3$ ), representing the signal-to-noise ratio where aging effects dominate over confounding biological variation. Two trajectories (Y\_308 and Y\_272) met these criteria and were selected for subsequent GeneSwitches and attractor landscape analyses.

**Figure 1:**
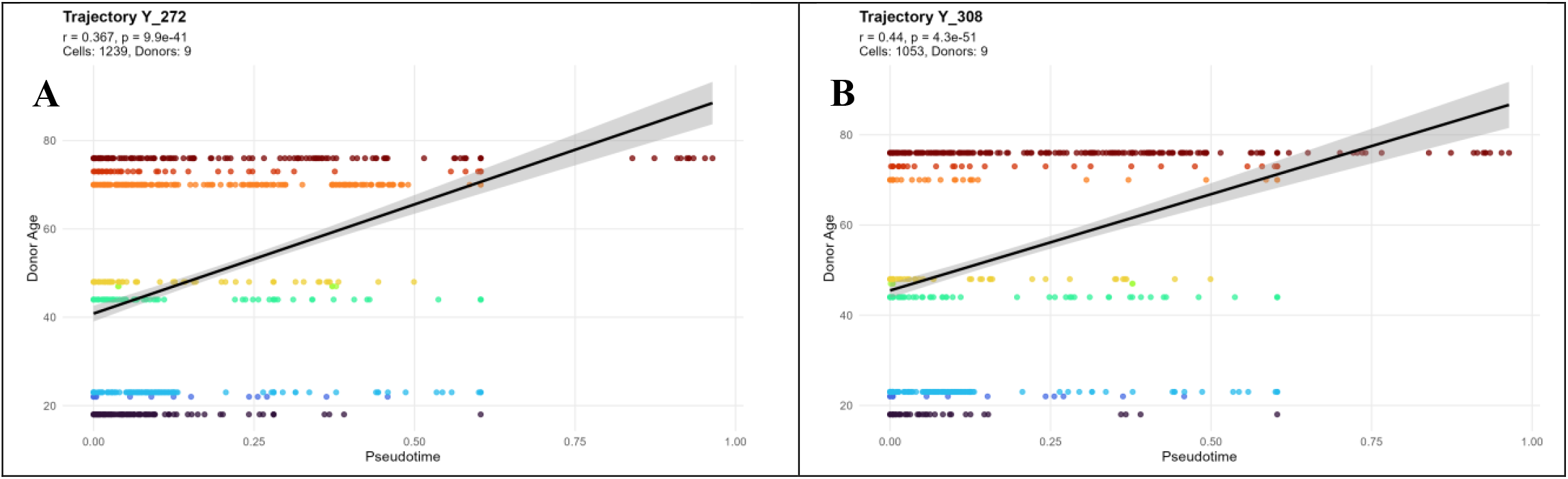
Trajectories Y_272 and Y_308. Trajectory Y_272 (panel A) shows significant correlation between pseudotime and donor chronological age (Spearman r = 0.367, p = 9.9 × 10−41, n = 1,239 cells). Trajectory Y_308 (panel B) demonstrates the strongest age-pseudotime correlation (Spearman r = 0.440, p = 4.3 × 10−51, n = 1,053 cells).

### Gene Switch Identification

GeneSwitches analysis (Cao et al., 2020) identified genes exhibiting switch-like expression transitions along each pseudotime trajectory, providing the molecular foundation for Boolean network construction. These switch genes represent regulatory factors whose binary expression states (ON/OFF) capture meaningful cellular transitions during aging, making them ideal candidates for Boolean modeling where gene expression is discretized to binary values.

For trajectory Y_272 (Figure 2a and Table 3), GeneSwitches identified 25 significant switch genes (FDR < 10^−7^), predominantly showing upregulation with aging (24 up, 1 down). The top-ranked switches included NFKBIA (FDR = 5.0 × 10^−24^, pseudo-*R*^2^ = 0.084, switch time = 0.38), OVOL1 (FDR = 7.7 × 10^−18^, pseudo-*R*^2^ = 0.058, switch time = 0.22), and NEDD4L (FDR = 5.9 × 10^−14^, pseudo-*R*^2^ = 0.046, switch time = 0.13). These genes exhibited logistic expression transitions with high model fit quality, indicating their expression patterns closely approximated ideal binary switches. The temporal distribution of switch points spanned the entire pseudotime range (0.04 to 0.93), with notable clustering in early (0.04-0.13) and mid-trajectory regions (0.30-0.40), suggesting coordinated regulatory modules activating at specific aging stages.

**Table 3:** Top Switch Genes Identified in Y_272 and Y_308 Trajectories.

| Y_272<br>Gene | Direction | FDR | Y_308<br>Gene | Direction | FDR |
| --- | --- | --- | --- | --- | --- |
| NFKBIA | up | $5.0 \times 10^{-24}$ | IGFBP3 | up | $4.1 \times 10^{-18}$ |
| OVOL1 | up | $7.7 \times 10^{-18}$ | SPON2 | up | $6.5 \times 10^{-12}$ |
| NEDD4L | up | $5.9 \times 10^{-14}$ | BGN | up | $1.3 \times 10^{-11}$ |
| NFKB1 | up | $5.9 \times 10^{-14}$ | SLC35F1 | down | $1.0 \times 10^{-10}$ |
| PHLDA2 | up | $5.9 \times 10^{-14}$ | SOX6 | up | $1.0 \times 10^{-10}$ |
| EPHA2 | up | $8.0 \times 10^{-14}$ | WNT4 | up | $5.8 \times 10^{-10}$ |
| ZC3H12A | up | $8.0 \times 10^{-14}$ | GAS6 | up | $4.3 \times 10^{-9}$ |
| LGALS7 | down | $2.3 \times 10^{-13}$ | TSC22D1 | down | $7.6 \times 10^{-9}$ |
| MGST1 | up | $1.8 \times 10^{-10}$ | PRR7 | up | $2.1 \times 10^{-8}$ |
| PANX1 | up | $1.8 \times 10^{-10}$ | GLUL | down | $3.5 \times 10^{-8}$ |

The ten most statistically significant switch genes for each validated aging trajectory, ranked by false discovery rate (FDR). Switch genes were identified by GeneSwitches analysis, which fits logistic regression models to binary expression patterns along pseudotime to detect genes exhibiting sigmoid-like transitions characteristic of cellular state changes. Direction indicates whether gene expression increases (up) or decreases (down) with advancing pseudotime. Y\_272 exhibited predominantly upregulated switches (24 of 25 total), while Y\_308 showed more balanced directionality (10 up, 3 down of 13 total). All listed genes passed the $FDR < 10^{-7}$ significance threshold.

**Figure 2:**
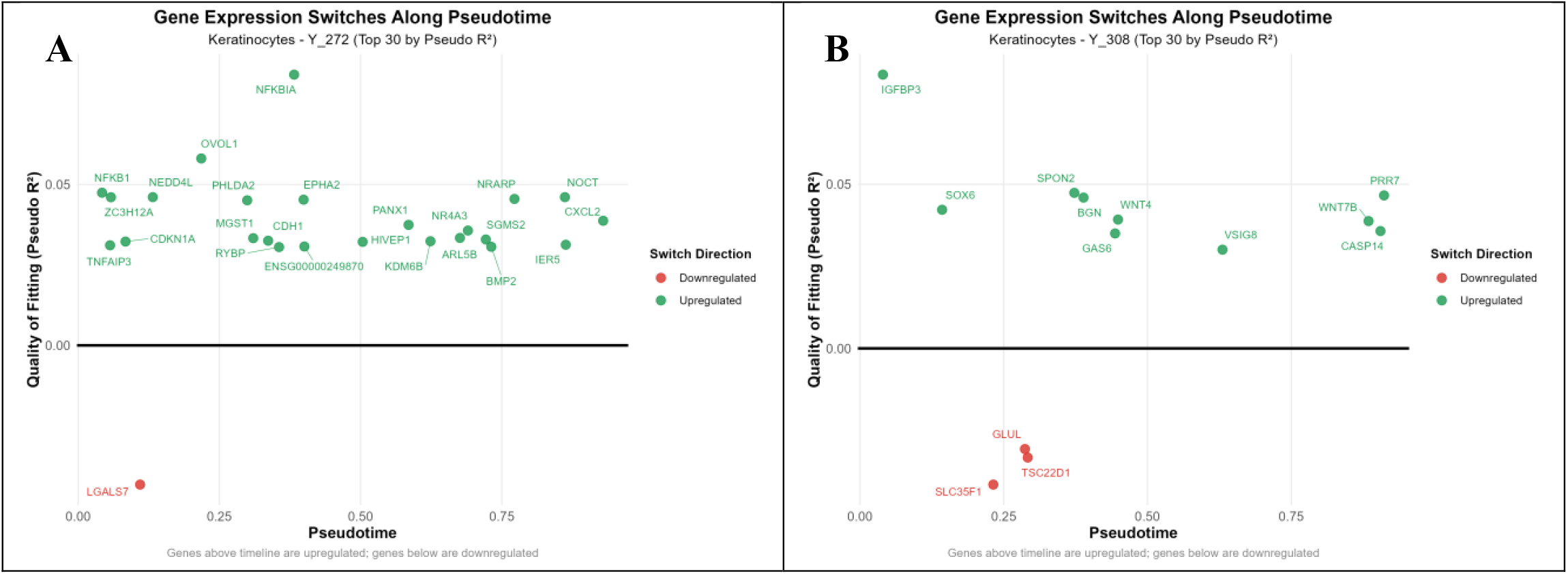
GeneSwitches Analysis. Analysis for Y_272 trajectory (panel A) shows temporal expression transitions of 25 significant switch genes along pseudotime, with color indicating upregulation (green) or downregulation (red). Analysis for Y_308 trajectory (panel B) displays temporal expression patterns of 13 significant switch genes, showing more balanced directionality (10 up, 3 down) compared to Y_272.

For trajectory Y_308 (Figure 2b and Table 3), GeneSwitches identified 13 significant switch genes (FDR < 10^−7^), showing more balanced directionality (10 up, 3 down). Top switches included IGFBP3 (FDR = 4.1 × 10^−18^, pseudo-*R*^2^ = 0.083, switch time = 0.04), SPON2 (FDR = 6.5 × 10^−12^, pseudo-*R*^2^ = 0.047, switch time = 0.37), and BGN (FDR = 1.3 × 10^−11^, pseudo-*R*^2^ = 0.046, switch time = 0.39). The earlier switch time of IGFBP3 (0.04) suggests it may serve as an initiating event in this aging trajectory, while the mid-to-late switches (0.23-0.91) indicate progressive regulatory changes throughout the aging process.

#### Gene Regulatory Network Construction

SCENIC analysis (Aibar et al., 2017) integrated GENIE3 random forest predictions (Huynh-Thu et al., 2019) with RcisTarget transcription factor binding motif enrichment (Aibar S and Aerts S, 2016) to construct gene regulatory networks linking switch genes to their potential regulators. For trajectory Y_272, SCENIC identified 1,638 regulatory edges connecting 134 transcription factors to 606 target genes, with overwhelming bias toward activation relationships (1,599 activating, 39 inhibiting). All edges received motif support from RcisTarget analysis, indicating high-confidence regulatory predictions grounded in transcription factor binding site enrichment. The network exhibited scale-free topology typical of biological regulatory networks, with a small number of transcription factors (hub regulators) controlling large numbers of target genes. For trajectory Y_308, SCENIC generated a slightly smaller but structurally similar network comprising 1,538 edges linking 131 transcription factors to 572 targets. This network showed comparable activation bias (1,451 activating, 87 inhibiting) with universal motif support. The reduced network size relative to Y_272 likely reflects the smaller number of switch genes identified in Y_308, propagating through the regulatory inference to produce a more compact network architecture.

### Boolean Network Construction

The SCENIC-derived gene regulatory networks were transformed into executable Boolean logic rules through a systematic template-based inference approach that tested logical combinations of regulators against empirical binary expression patterns. Rule construction proceeded by binarizing gene expression along pseudotime-ordered cells, generating candidate Boolean rules using templates (single activator, activator OR, activator AND, repressor variants), and scoring each rule based on prediction accuracy against the binary expression data.

For trajectory Y_272, the Boolean network comprised 613 genes with mean rule quality of 0.673 (median = 0.676). Template-based methods successfully generated 606 rules (99.0% of network), with only 7 genes requiring fallback self-activation rules due to insufficient regulatory evidence. The network exhibited moderate complexity with mean 2.54 regulators per gene, skewed toward simpler architectures: 7 genes with zero regulators (constant genes), 66 with single regulators, 126 with two regulators, and 414 with three or more regulators. Logical operators showed strong bias toward OR logic (531 rules) over AND logic (0 rules), with minimal use of NOT operators (3 rules), reflecting SCENIC’s predominant identification of activation relationships.

For trajectory Y_308, the Boolean network contained 575 genes with nearly identical mean rule quality (0.674, median = 0.675), suggesting consistent rule inference performance across trajectories. Template-based methods generated 572 rules (99.5%), with only 3 genes requiring fallback rules. Network complexity closely paralleled Y_272 (mean 2.53 regulators per gene), with similar architectural distribution: 3 zero-regulator genes, 67 single-regulator genes, 126 two-regulator genes, and 379 with three or more regulators. Logical operator distribution again favored OR logic (491 rules) over AND (0 rules), with slightly more NOT operators (5 rules) than Y_272.

The high-quality rules for both networks achieved scores clustering tightly around 0.68, indicating reliable prediction of gene states from regulator combinations. Top-scoring rules included NFKBIA, CLOCK, and NFKB1 for Y_272 (scores 0.682, 0.682, 0.682) and TEAD1, BACH2, and RBPJ for Y_308 (scores 0.683, 0.682, 0.681). No rules exceeded the stringent 0.75 quality threshold, reflecting the inherent biological noise in single-cell data and the challenge of perfectly predicting stochastic gene expression with deterministic Boolean logic. Nevertheless, the consistent 67% prediction accuracy across both networks provided sufficient signal for meaningful attractor analysis.

### Attractor Landscape Analysis

Synchronous Boolean network simulation of the Y_272 network (613 genes) using random state sampling identified a single dominant attractor that captured all sampled initial states, indicating universal convergence behavior. This attractor manifested as a 20-state limit cycle rather than a point attractor, representing a dynamic oscillation among 20 related gene expression configurations that the network circulates through in a deterministic sequence. The attractor’s aging score of +4.362 (measured by projection onto the young-old reference axis) placed it firmly in the aged regime, exceeding the baseline network aging score by a factor of two. The complete dominance of this single aged attractor (basin size = 100%) reveals the fundamental character of the Y_272 aging program: aging represents convergence toward a stable aged configuration that acts as a universal attractor basin in gene expression state space. Regardless of initial conditions, the regulatory logic encoded in the Boolean rules drives the system toward this aged attractor, suggesting that for trajectory Y_272 aging involves establishing a robust regulatory architecture that stabilizes aged gene expression patterns. The attractor’s cyclic nature indicates that even in the aged configuration, the network exhibits dynamic behavior, cycling through related expression states rather than settling into complete stasis. Entropy analysis of the attractor revealed zero Shannon entropy, indicating perfectly deterministic transitions among the 20 attractor states with no probabilistic variation. The absence of alternative attractors or complex loose attractors indicates that the Y_272 regulatory architecture, once perturbed from the aged attractor, rapidly returns to it, exhibiting strong homeostatic maintenance of the aged state. The composite aging contribution metric, which weights each attractor’s aging score by its basin size, yielded a value of +2.181 for the single attractor. This value represents the network’s overall bias toward aged configurations, quantifying the probability-weighted aging phenotype that emerges from the attractor landscape.

In contrast to trajectory Y_272, Boolean network simulation of trajectory Y_308 (575 genes) identified a complex six-attractor landscape comprising three point attractors (single-state steady states) and three cycle attractors (18-state and 36-state cycles). This architectural diversity indicates that cells along trajectory Y_308 can settle into multiple distinct stable configurations rather than converging universally to a single state. The attractor landscape exhibited extreme dominance by a single youthful attractor that captured 95.8% of all initial states, representing an overwhelming basin of attraction. This dominant attractor manifested as an 18-state cycle with an aging score of −0.538, placing it firmly in the youthful regime. The remaining five attractors shared the remaining 4.2% of state space, with individual basin sizes ranging from 0.48% to 1.6%. All six attractors showed negative (youthful) aging scores ranging from −0.485 to −0.538, indicating the entire attractor landscape biases toward youthful gene expression configurations. This attractor architecture reveals a fundamentally different aging mechanism from trajectory Y_272. Rather than convergence toward an aged state, trajectory Y_308 aging represents progressive failure to maintain residence in the dominant youthful attractor. The large basin of attraction surrounding the youthful state suggests strong homeostatic mechanisms that actively maintain youthful configurations in young cells. Aging in this model corresponds to perturbations or regulatory changes that shift cells out of the dominant youthful basin into the smaller alternative basins, representing loss of youthful regulatory control rather than gain of aged regulatory programs. Stability analysis revealed that five of the six attractors achieved perfect stability (value = 1.0), with only the sixth attractor showing slight deviation (0.996) due to minimal entropy (0.011). The weighted aging score calculation integrated all six attractors’ contributions, producing an overall network aging score of −0.536. This negative value reflects the overwhelming dominance of youthful attractors in the landscape, with the large basin of the dominant attractor (−0.515 composite contribution) far outweighing the smaller contributions of the alternative attractors (ranging from −0.002 to −0.008).

### Perturbation Analysis and Therapeutic Targets

Systematic perturbation analysis tested all 613 genes under knockdown and overexpression conditions, plus combinatorial double perturbations and mixed knockdown/overexpression pairs for the top single-gene targets. BACH2 knockdown emerged as the dominant rejuvenation intervention, achieving a network aging score of −1.565 (Δ= −3.746 from baseline +2.181, representing 98.9% improvement on the scale to maximal observed improvement; see Table 4).

**Table 4:** Top Therapeutic Targets for Trajectory Y_272 Ranked by Aging Score Improvement.

| Rank | Target | Type | Aging Score | $\Delta$ |
| --- | --- | --- | --- | --- |
| 1 | BACH2 + ARL5B | Double KD | −1.608 | −3.789 |
| 2 | BACH2 + ATF6 | Double KD | −1.605 | −3.786 |
| 3 | BACH2 + ATF1 | Double KD | −1.597 | −3.778 |
| 4 | BACH2 + ADNP | Double KD | −1.588 | −3.769 |
| 5 | BACH2 | Single KD | −1.565 | −3.746 |
| 6 | BACH2 (KD) + ABCF2 (OE) | Mixed | −1.565 | −3.746 |
| 7 | BACH2 (KD) + ABL1 (OE) | Mixed | −1.565 | −3.746 |
| 8 | BACH2 (KD) + ACAA1 (OE) | Mixed | −1.565 | −3.746 |
| 9 | BACH2 (KD) + ACO1 (OE) | Mixed | −1.565 | −3.746 |
| 10 | BACH2 (KD) + ADARB1 (OE) | Mixed | −1.565 | −3.746 |

Systematic perturbation analysis identifying interventions that shift the Y\_272 attractor landscape toward more youthful configurations. Aging score represents the projection of attractor states onto the young-old reference axis (positive = aged, negative = youthful). Delta indicates improvement from the baseline network aging score of +2.181. BACH2 knockdown emerged as the dominant single-gene target, with double knockdown combinations providing modest additional benefit. Mixed perturbations (knockdown + overexpression) showed no synergistic enhancement beyond single BACH2 knockdown. KD = knockdown; OE = overexpression.

**Table 5:** Top Therapeutic Targets for Trajectory Y_308 Ranked by Aging Score Improvement.

| Rank | Target | Type | Aging Score | $\Delta$ |
| --- | --- | --- | --- | --- |
| 1 | ASCL2 + ATF6 | Double KD | -0.623 | -0.088 |
| 2 | ASCL2 + AFF4 | Double KD | -0.617 | -0.081 |
| 3 | ASCL2 + BACH1 | Double KD | -0.611 | -0.075 |
| 4 | ASCL2 + ACAA1 | Double KD | -0.610 | -0.075 |
| 5 | ASCL2 (KD) + ATF3 (OE) | Mixed | -0.609 | -0.073 |
| 6 | ASCL2 (KD) + BGN (OE) | Mixed | -0.608 | -0.073 |
| 7 | ASCL2 | Single KD | -0.608 | -0.072 |
| 8 | ASCL2 (KD) + CNOT4 (OE) | Mixed | -0.608 | -0.072 |
| 9 | ASCL2 (KD) + CERS3 (OE) | Mixed | -0.607 | -0.071 |
| 10 | ASCL2 (KD) + AHCTF1 (OE) | Mixed | -0.606 | -0.070 |

Perturbation analysis results for trajectory Y\_308, which exhibited a fundamentally different attractor architecture from Y\_272. The baseline network aging score of -0.536 (already youthful) reflects the dominance of a youthful attractor capturing 95.8% of the state space. ASCL2 knockdown emerged as the primary intervention target, with the ASCL2 + ATF6 double knockdown combination demonstrating synergistic enhancement (Delta = -0.088 vs. -0.072 for single knockdown). The smaller magnitude of improvements compared to Y\_272 reflects the already youthful baseline configuration of this trajectory's attractor landscape. KD = knockdown; OE = overexpression.

BACH2 knockdown fundamentally restructured the attractor landscape from a single aged attractor to a configuration biased toward youthful states. The magnitude of the aging score shift (−3.746) exceeded the original baseline aging score (+2.181) by 72%, indicating that BACH2 perturbation not only neutralized the aged attractor but drove the network substantially into youthful territory. This dramatic effect positions BACH2 as a master regulator of the Y_272 aging program, whose activity appears necessary for maintaining the aged attractor’s dominance. None of the switch genes identified by GeneSwitches analysis provided beneficial perturbation targets in Y_272. All 25 switch gene knockdowns and overexpressions yielded either no change or increased aging scores, revealing a critical distinction between genes whose expression changes during aging (switch genes) and genes whose perturbation can reverse aging (therapeutic targets). The switch genes, while marking the aging trajectory, appear to be passengers rather than drivers of the aged attractor’s stability. BACH2, though not identified as a switch gene, emerged from SCENIC as a key transcription factor in the regulatory network, suggesting it operates as a regulatory hub rather than a trajectory marker.

Double perturbation analysis tested combinations of the top five single-gene targets (BACH2, ARL5B, ATF6, ATF1, ADNP) in all possible knockdown pairs, overexpression pairs, and mixed configurations. The top four combinations all involved BACH2 knockdown paired with knockdown of a second target: BACH2 + ARL5B (Δ= −3.789, 100% improvement), BACH2 + ATF6 (Δ= −3.786, 99.9%), BACH2 + ATF1 (Δ= −3.778, 99.7%), and BACH2 + ADNP (Δ= −3.769, 99.5%). The modest additional improvements over BACH2 alone (ranging from 0.043 to 0.001 units) indicate largely independent effects rather than synergistic interactions, with BACH2’s dominant effect difficult to enhance through combinatorial approaches.

The perturbation results were consistent across intervention modes, with knockdown producing identical effects to mixed knockdown/overexpression for all tested combinations. Conversely, the worst-performing perturbations involved knockdown of genes that stabilize youthful configurations. ARL5B knockdown produced the most severe pro-aging effect (Δ= +2.139, representing movement away from youthful states), followed by combinations of ARL5B with other targets. This bidirectional response spectrum (BACH2 knockdown improves aging, ARL5B knockdown worsens it) demonstrates that the Boolean network captures both protective and deleterious regulatory factors, providing a complete map of the aging regulatory landscape.

Perturbation analysis of the Y_308 network identified ASCL2 knockdown as the most effective single-gene intervention, achieving a network aging score of −0.608 (Δ= −0.072 from baseline −0.536, representing 82.1% improvement toward maximal observed change). While this effect size appears modest compared to Y_272’s BACH2 results, it must be interpreted in the context of Y_308’s distinct baseline: the network already exhibits strong youthful bias (−0.536), leaving less dynamic range for improvement than Y_272’s aged baseline (+2.181). ASCL2 knockdown’s mechanism likely involves reinforcing the dominant youthful attractor’s basin of attraction, making it even more difficult for cells to escape into the alternative aged-biased basins. The relatively small magnitude of improvement suggests that Y_308’s aging process represents subtle erosion of youthful attractor dominance rather than wholesale transition to aged states, consistent with the “aging as departure” model where most cells remain in youthful configurations even during chronological aging. Double perturbation analysis revealed more substantial synergistic effects in Y_308 compared to Y_272. The top combination, ASCL2 + ATF6 double knockdown, achieved Δ= −0.088 (100% improvement), representing a 22% enhancement over ASCL2 alone. This synergy suggests that ASCL2 and ATF6 operate in partially independent pathways that converge to maintain youthful attractor dominance, and simultaneous perturbation of both pathways produces additive or synergistic effects. The next-best combinations ASCL2 + AFF4 (Δ= −0.081, 92.4%) and ASCL2 + BACH1 (Δ= −0.075, 85.7%) showed progressively decreasing synergy, indicating a hierarchy of pathway interactions.

Mixed knockdown/overexpression combinations also showed promise, with ASCL2 knockdown + ATF3 overexpression achieving Δ= −0.073 (83.6% improvement). This near-equivalence to ASCL2 knockdown alone suggests that ATF3 overexpression provides minimal additional benefit when combined with ASCL2 perturbation, possibly because both interventions converge on the same regulatory module. In contrast to Y_272, Y_308 showed minimal effects from switch gene perturbations. The top switch genes identified by GeneSwitches (IGFBP3, SPON2, BGN, SOX6) yielded small or non-significant aging score changes when perturbed, consistent with the pattern observed in Y_272 where switch genes mark aging trajectories but do not necessarily control attractor landscape architecture. However, some switch genes did show modest effects: BGN knockdown achieved Δ= −0.061 (ranked 22nd), while PRR7 overexpression showed minimal benefit (Δ= −0.002). The perturbation spectrum in Y_308 proved narrower than Y_272, with the best improvement (Δ= −0.088) and worst deterioration (VSIG8 knockdown, Δ= +0.003) spanning only 0.091 units compared to Y_272’s range of 5.928 units (from −3.789 to +2.139). This compressed dynamic range reflects Y_308’s baseline proximity to optimal youthful configuration, where even the best interventions can only marginally improve an already favorable attractor landscape, while deleterious perturbations struggle to substantially destabilize the dominant youthful basin.

## Discussion

Two distinct aging trajectories were identified in human skin keratinocytes through the application of the PRISM pipeline, each characterized by fundamentally different regulatory architectures and attractor landscape dynamics. Trajectory Y_272 was found to represent “aging as convergence,” wherein cells were driven toward a single dominant aged attractor state stabilized by stress-response transcription factor networks. In contrast, trajectory Y_308 was characterized as “aging as departure,” where cells progressively escaped from a dominant youthful attractor basin through accumulation of senescence-associated secretory factors. A critical distinction was revealed between genes exhibiting age-related expression changes (phenotypic markers) and genes controlling attractor landscape architecture (regulatory controllers). Switch genes identified through GeneSwitches analysis marked the aging trajectories but were found to be largely ineffective as perturbation targets for reversing aging scores. Instead, master regulators operating at higher levels of the regulatory hierarchy were identified as the most promising intervention targets: BACH2 for trajectory Y_272 and ASCL2 for trajectory Y_308.

The phenotypic signature of trajectory Y_272 was characterized by coordinated upregulation of genes involved in terminal differentiation, barrier reinforcement, and chronic stress adaptation. NFKBIA, the canonical negative feedback inhibitor of NF-κB, was identified as a top switch gene, suggesting that keratinocytes along this trajectory enter a regime of sustained low-grade inflammation with compensatory feedback control. This pattern is consistent with the “inflammaging” phenomenon, where chronic activation of inflammatory pathways contributes to age-related dysfunction (Franceschi & Campisi, 2014). Additional switch genes including OVOL1 (a master regulator of epidermal differentiation and barrier maintenance) and NEDD4L (an E3 ubiquitin ligase that antagonizes TGF-*β* signaling) indicated that Y_272 cells adopt a terminally differentiated, barrier-reinforced phenotype. This configuration may represent a compensatory response to chronic environmental damage rather than simple cellular deterioration. Despite these visible phenotypic changes, BACH2 was identified as the dominant controller of the aged attractor state through perturbation analysis. BACH2 knockdown produced a dramatic shift in aging score (Δ= −3.746), collapsing the aged attractor and driving the network toward youthful configurations. This finding positions BACH2 as a transcriptional “lock” that actively maintains the chronic stress state. The co-perturbation partners of BACH2 (ATF6, ATF1, ARL5B, and ADNP) revealed a multi-layered regulatory infrastructure involving ER stress sensing, metabolic regulation, nutrient sensing, and chromatin remodeling, all converging to stabilize the aged attractor.

A contrasting aging mechanism was observed in trajectory Y_308, where cells drifted away from a dominant youthful attractor through accumulation of senescence-associated secretory phenotype (SASP) factors and extracellular matrix (ECM) remodeling proteins. IGFBP3, a canonical SASP factor (Özcan et al., 2016), was identified as the earliest switch gene (pseudotime = 0.04), suggesting it may serve as an initiating event in this aging trajectory. Additional switch genes including SPON2 (spondin-2) and BGN (biglycan) indicated active ECM remodeling coupled to immune system communication through damage-associated molecular pattern (DAMP) signaling (Moreth et al., 2012). The attractor landscape of Y_308 was characterized by a dominant youthful attractor capturing 95.8% of initial states, with five smaller alternative attractors sharing the remaining state space. All attractors exhibited negative (youthful) aging scores, indicating that the regulatory architecture inherently biases toward youthful configurations. Aging in this model was interpreted as progressive failure to maintain residence in the dominant youthful basin rather than convergence toward an aged state. ASCL2 knockdown was identified as the most effective single-gene intervention for Y_308 (Δ= −0.072, 82.1% improvement). The mechanism was interpreted as reinforcement of the dominant youthful attractor’s basin of attraction, making escape into alternative basins more difficult. Notably, synergistic effects were observed with double perturbations in Y_308, with ASCL2 + ATF6 knockdown achieving 22% enhancement over ASCL2 alone, suggesting that multiple partially independent pathways converge to maintain youthful attractor dominance.

One of the most significant findings of this analysis was the systematic divergence between genes marking aging trajectories and genes controlling attractor landscape architecture. All 25 switch genes identified in Y_272 and all 13 switch genes in Y_308 proved ineffective or detrimental when perturbed, yielding either no change or increased aging scores. This observation challenges conventional approaches to aging research that focus primarily on differential gene expression as the basis for therapeutic target identification. The switch genes, while exhibiting the clearest age-related expression changes, appear to function as downstream effectors or passengers rather than upstream drivers of the aging process. In contrast, master regulators such as BACH2 and ASCL2 operate at higher levels of the regulatory hierarchy, controlling the stability and accessibility of attractor basins without necessarily exhibiting dramatic expression changes themselves. This distinction has important implications for therapeutic development, suggesting that expression-based biomarkers of aging may be poor guides for identifying intervention targets.

The present analysis was conducted on the same Human Cell Atlas eyelid skin dataset originally characterized by Zou et al. (2021). Both analyses converge on similar biological themes, including chronic NF-κB-driven inflammation, loss of youthful basal keratinocyte programs, ECM disorganization, and IGFBP3-linked changes in basal compartments. However, several key differences in interpretation and target identification emerged. In their original analysis, Zou et al. emphasized fibroblasts as the most age-vulnerable lineage and identified HES1 as a fibroblast geroprotective transcription factor and KLF6 as protective in keratinocytes, proposing quercetin as a protective compound through restoration of these factors. The present PRISM analysis, in contrast, identified keratinocyte-specific aging mechanisms and different master regulators (BACH2, ASCL2) whose perturbation was predicted to reshape the attractor landscape and reverse aging scores. Additionally, Zou et al. described aging primarily as transcriptional drift and pathway-level decline mapped onto a single differentiation trajectory. The PRISM analysis distinguished two mechanistically distinct aging modes (convergence versus departure) with different attractor architectures and different optimal intervention strategies. The identification of trajectory-specific targets suggests that personalized approaches to skin rejuvenation may be necessary depending on which aging mechanism predominates in a given context.

Several limitations of the present analysis should be acknowledged. First, Boolean network models necessarily discretize gene expression to binary states, which may not capture dosage-dependent effects or continuous gradients of gene activity. The consistent 67% prediction accuracy of Boolean rules across both networks, while sufficient for meaningful attractor analysis, reflects the inherent biological noise in single-cell data and the challenge of perfectly predicting stochastic gene expression with deterministic logic. Second, the perturbation analysis was conducted computationally by fixing gene states to 0 (knockdown) or 1 (overexpression), which represents an idealized intervention that may not be achievable in experimental settings. Partial knockdowns, incomplete overexpression, and off-target effects in real biological systems may produce different results than those predicted by the Boolean model. Third, the analysis was limited to keratinocytes due to their dominance in the dataset (80.4% of cells) and successful trajectory validation. Other cell types, including fibroblasts that Zou et al. previously identified as highly age-vulnerable, were not analyzed due to insufficient cell numbers or failure to meet the age-correlation threshold for trajectory validation. The aging mechanisms identified here may therefore represent only a subset of the total aging program operating in human skin. Fourth, the temporal resolution of the dataset (cross-sectional sampling from donors of different ages) limits the ability to distinguish true longitudinal aging trajectories from population heterogeneity. The pseudotime ordering assumes that cells from older donors occupy later positions along a continuous aging trajectory, which may not fully capture the complexity of individual aging histories. Finally, the regulatory network construction relied on SCENIC’s inference of transcription factor-target relationships from co-expression patterns, which may include false positive edges and miss true regulatory relationships not captured by the algorithm’s assumptions. The Boolean rules derived from this network inherit any inaccuracies in the underlying regulatory structure.

The application of Boolean network modeling to aging keratinocyte trajectories revealed two mechanistically distinct aging modes operating in human skin: stress-adapted convergence to aged attractor states (Y_272) and SASP-driven departure from youthful attractor basins (Y_308). A critical distinction was identified between phenotypic markers of aging, which exhibit clear expression changes but prove ineffective as intervention targets, and regulatory controllers that govern attractor landscape architecture and represent promising candidates for cellular rejuvenation. BACH2 was identified as the master regulator of the Y_272 aged attractor, with knockdown predicted to produce dramatic reversal of aging scores. ASCL2 was identified as the top target for Y_308, with synergistic enhancement observed through combinatorial perturbation with ATF6. These findings provide concrete therapeutic hypotheses for experimental validation and demonstrate the utility of dynamical systems approaches for moving beyond description of aging to mechanistic understanding and rational intervention design. More broadly, the PRISM framework illustrates how integration of trajectory analysis, gene regulatory network inference, Boolean modeling, and systematic perturbation testing can transform static transcriptomic snapshots into actionable predictions about cellular control. By identifying not just what changes with age but what controls those changes, this approach opens new avenues for developing targeted strategies to reverse cellular aging.

## Materials and Methods

### Data Acquisition and Preprocessing

Single-cell RNA sequencing (scRNA-seq) data were obtained from 10X Genomics CellRanger output files. Raw count matrices from multiple biological donors were loaded into R and processed using Seurat v5. Doublet detection was performed using scDblFinder (Germain et al., 2021) to identify and remove artifactual cell multiplets prior to downstream analysis. Quality control filters were applied based on mitochondrial gene percentage (cells exceeding 20% mitochondrial reads were excluded) and feature counts to remove low-quality cells and potential empty droplets. Following quality control, individual donor datasets were merged into a single Seurat object. The merged dataset was subjected to standard preprocessing including log-normalization, identification of highly variable features, and scaling. Principal component analysis (PCA) was performed for initial dimensionality reduction, followed by Uniform Manifold Approximation and Projection (UMAP) embedding for visualization. To address batch effects arising from donor-to-donor technical variation while preserving biological signal, donor-level integration was performed using canonical correlation analysis (CCA).

### Cell Type Annotation and Donor Normalization

Automated cell type annotation was performed using SingleR (Aran et al., 2019) with the Human Primary Cell Atlas reference database. To prevent donor-specific biases from confounding downstream pseudotime and attractor analyses, a donor normalization strategy was implemented to ensure balanced donor representation within each cell type. Normalization was achieved through random downsampling to match the minimum donor count within each cell type, ensuring perfect balance across all donors. Cell types that failed to achieve balanced representation across all donors were excluded from further analysis. For each cell type passing the normalization criteria, a separate Seurat object was created and subjected to cell-type-specific dimensionality reduction (PCA and UMAP). Summary statistics documenting pre- and post-normalization donor distributions were generated, along with frequency tables recording the total number of cells per cell type after filtering.

### Pseudotime Trajectory Inference

Cell-type-specific Seurat objects were converted to Monocle3 CellDataSet format for trajectory inference. Pseudotime analysis was rooted from cells belonging to the youngest donor, representing the earliest cellular state in the aging process. All trajectory branches emanating from this root were identified, and each branch was validated by calculating Spearman correlation coefficients between pseudotime values and donor chronological age. A conservative correlation threshold (*r* ≥ 0.3, *p* < 0.05) was established as the minimum signal-to-noise ratio at which aging effects dominate over non-aging factors in the trajectory. Only branches exceeding this threshold were retained for downstream analysis. For each validated branch, a separate CellDataSet containing only cells along that trajectory path was saved, along with age-versus-pseudotime visualizations and comprehensive branch statistics documenting correlations, *p*-values, cell counts, and donor representation.

### Gene Switch Identification

The GeneSwitches package (Cao, et al., 2020) was applied to identify genes exhibiting switch-like expression transitions along validated pseudotime trajectories. Raw count data were converted to binary expression matrices using mixture model binarization, by which expression thresholds were adaptively determined for each gene. Fixed cutoff binarization was employed as a fallback when mixture models failed to converge. Each gene’s expression was modeled as a logistic function of pseudotime, with parameters estimated including switch timing (the pseudotime coordinate at which the transition occurs), direction (upregulation or downregulation), and fit quality (pseudo-*R*^2^). Results were filtered by false discovery rate (FDR < 0.05) to retain only statistically significant switch genes. Output included gene identifiers, switch directions, FDR values, pseudotime coordinates, and model fit statistics, providing the molecular foundation for subsequent Boolean network construction.

### Gene Regulatory Network Construction

Gene regulatory networks (GRNs) were constructed by integrating SCENIC’s GENIE3 random forest predictions (Huynh-Thu et al., 2019) with RcisTarget transcription factor binding motif enrichment analysis. Potential regulatory interactions were predicted based on co-expression patterns across cells ordered by pseudotime using GENIE3. These predictions were validated using RcisTarget to confirm transcription factor binding motif enrichment in target gene promoter regions, ensuring biologically grounded regulatory relationships. The resulting network was filtered to focus on relationships involving switch genes identified in the previous step. Binarized expression patterns between regulators and targets were correlated to quantify relationship strength. SCENIC metadata, including regulatory direction (activation versus repression) and motif confidence scores, were incorporated into the final edge list. The complete regulatory network served as input for Boolean rule inference.

### Boolean Network Inference

The SCENIC-derived gene regulatory network was transformed into executable Boolean logic rules through a template-based inference approach. Gene names were sanitized for BoolNet compatibility, and optimal regulators for each target gene were selected using SCENIC metadata (correlation strength, regulatory direction, motif confidence). A multi-method inference strategy was employed in which logical combinations of AND, OR, and NOT operators with up to *k* regulators were exhaustively tested to identify rules that best predicted target gene binary expression patterns along pseudotime-ordered cells. Each candidate rule was assigned a quality score based on prediction accuracy, consistency with SCENIC-indicated regulatory signs, and motif evidence. For genes lacking strong regulatory evidence, intelligent fallbacks including self-activation rules were implemented to ensure complete network coverage. All rules were validated for BoolNet syntax compatibility. The final output included a detailed rule analysis table, a flat regulator table, and the complete rule set object for loading into the BoolNet framework.

### Attractor Identification

Boolean networks were loaded into BoolNet’s synchronous dynamics framework for attractor identification (Müssel et al., 2010). For networks containing 29 or fewer genes, exhaustive state space exploration was performed, guaranteeing identification of all attractors. For larger networks, adaptive random sampling was employed, with sample size scaled according to network complexity. Attractors were classified as either point attractors (steady states where no genes change between time steps) or cycle attractors (deterministic oscillations among multiple gene expression configurations). For each attractor, basin of attraction statistics were computed, indicating the fraction of possible initial states that eventually reach that attractor. These statistics provided measures of attractor stability and dominance within the network’s state space.

### Aging Score Computation

Aging scores were computed for each attractor by projecting attractor binary gene expression patterns onto a young-old reference axis. This reference axis was derived from the mean binarized expression profiles of the 20% youngest and 20% oldest cells along the pseudotime trajectory. The scalar projection of each attractor state onto this axis quantified how “young-like” (negative scores) or “old-like” (positive scores) each attractor appeared. Attractor stability was assessed through Shannon entropy analysis of state transitions. An overall network aging score was computed by weighting individual attractor aging scores by their respective basin sizes, representing the probability-weighted aging bias of the entire attractor landscape.

### Perturbation Analysis

Systematic perturbation analysis was performed to identify gene targets capable of shifting the attractor landscape toward more youthful configurations. Single-gene perturbations were tested by computationally fixing each gene to 0 (knockdown) or 1 (overexpression), followed by recalculation of the network’s attractor landscape and aging score for each perturbation. The change in aging score (Δ) relative to the unperturbed baseline was used to quantify each intervention’s rejuvenation potential. All perturbation results were ranked by aging score reduction. Comprehensive target tables were generated combining knockdown and overexpression results, with pivot tables indicating the optimal intervention mode (knockdown versus overexpression) for each gene. Double-gene perturbation analysis extended this approach to test combinations of top-ranked single-gene targets. All possible pairings were evaluated in three configurations: double knockdown (both genes fixed to 0), double overexpression (both genes fixed to 1), and mixed interventions (one gene knocked down, one overexpressed). Synergistic interactions, where combined perturbations produced greater rejuvenation effects than either single perturbation alone, were identified and integrated into a unified ranking system.

### Software and Statistical Analysis

All analyses were performed in R (version 4.4.1). Key packages included Seurat v5 (Hao et al., 2024) for scRNA-seq preprocessing and integration, Monocle3 (Trapnell et al., 2014; Cao et al., 2019) for trajectory inference, SingleR (Aran et al., 2019) for cell type annotation, GeneSwitches (Cao et al., 2020) for switch gene identification, SCENIC (Aibar et al., 2017) comprising GENIE3 and RcisTarget for gene regulatory network inference (Aibar S and Aerts S 2016), and BoolNet (Müssel et al., 2010) for Boolean network simulation and attractor analysis. Doublet detection was performed using scDblFinder (Germain et al., 2021). Statistical significance was assessed using Spearman correlation for pseudotime-age relationships and FDR correction for multiple testing in switch gene identification. Visualization was performed using ggplot2.

## Data and Code Availability

Processed data and analysis results are available via Zenodo at https://doi.org/10.5281/zenodo.17730145. The PRISM analysis pipeline source code is available at https://github.com/maypop-labs/prism. Additional information may be found at the project homepage: https://maypoplabs.org/projects/prism/

## References

Aibar S and Aerts S (2016). “RcisTarget.”

Aibar, S., González-Blas, C. B., Moerman, T., Huynh-Thu, V. A., Imrichova, H., Hulselmans, G., Rambow, F., Marine, J. C., Geurts, P., Aerts, J., van den Oord, J., Atak, Z. K., Wouters, J., & Aerts, S. (2017). SCENIC: single-cell regulatory network inference and clustering. Nature Methods, 14(11), 1083–1086. 10.1038/nmeth.4463

Aran, D., Looney, A. P., Liu, L., Wu, E., Fong, V., Hsu, A., Chak, S., Naikawadi, R. P., Wolters, P. J., Abate, A. R., Butte, A. J., & Bhattacharya, M. (2019). Reference-based analysis of lung single-cell sequencing reveals a transitional profibrotic macrophage. Nature Immunology, 20(2), 163–172. 10.1038/s41590-018-0276-y

Cao, J., Spielmann, M., Qiu, X., Huang, X., Ibrahim, D. M., Hill, A. J., Zhang, F., Mundlos, S., Christiansen, L., Steemers, F. J., Trapnell, C., & Shendure, J. (2019). The single-cell transcriptional landscape of mammalian organogenesis. Nature, 566(7745), 496–502. 10.1038/s41586-019-0969-x

Cao, Elaine Y., John F Ouyang, and Owen J L Rackham. “GeneSwitches: ordering gene expression and functional events in single-cell experiments.” Bioinformatics (Oxford, England) vol. 36, 10 (2020): 3273–3275. doi:10.1093/bioinformatics/btaa099

Franceschi, C., & Campisi, J. (2014). Chronic inflammation (inflammaging) and its potential contribution to age-associated diseases. The Journals of Gerontology Series A: Biological Sciences and Medical Sciences, 69(Suppl 1), S4–S9. 10.1093/gerona/glu057

Germain, P. L., Lun, A., Mace, A., & Robinson, M. D. (2021). Doublet identification in single-cell sequencing data using scDblFinder. F1000Research, 10, 979. 10.12688/f1000research.73600.1

Hao, Y., Stuart, T., Kowalski, M. H., Choudhary, S., Hoffman, P., Hartman, A., Srivastava, A., Mez, G., Longo, E., Karber, C., Radford, J., Zizic, H., Yadav, A., & Satija, R. (2024). Dictionary learning for integrative, multimodal and scalable single-cell analysis. Nature Biotechnology, 42(2), 293–304. 10.1038/s41587-023-01767-y

Huynh-Thu V. A., Irrthum A., Wehenkel L., and Geurts P. (2019). Inferring regulatory networks from expression data using tree-based methods. PLoS ONE, 5(9):e12776

Kuratomi, G., Komuro, A., Goto, K., Shinozaki, M., Miyazawa, K., Miyazono, K., & Imamura, T. (2005). NEDD4-2 (neural precursor cell expressed, developmentally down-regulated 4-2) negatively regulates TGF-β (transforming growth factor-β) signalling by inducing ubiquitin-mediated degradation of Smad2 and TGF-β type I receptor. Biochemical Journal, 386(3), 461–470.

Moreth, K., Iozzo, R. V., & Schaefer, L. (2012). Small leucine-rich proteoglycans orchestrate receptor crosstalk during inflammation. Cell Cycle, 11(11), 2084–2091.

Müssel, C., Hopfensitz, M., & Kestler, H. A. (2010). BoolNet—an R package for generation, reconstruction and analysis of Boolean networks. Bioinformatics, 26(10), 1378–1380. 10.1093/bioinformatics/btq124

Oeckinghaus, A., Hayden, M. S., & Ghosh, S. (2011). Crosstalk in NF-κB signaling pathways. Nature Immunology, 12(8), 695–708.

Ostapcuk, V., Mohn, F., Carl, S. H., Basters, A., Hess, D., Iesmantavicius, V., Lampersberger, L., Flemr, M., Pandey, A., Thoma, N. H., Betschinger, J., & Buhler, M. (2018). Activity-dependent neuroprotective protein recruits HP1 and CHD4 to control lineage-specifying genes. Nature, 557(7707), 739–743.

Özcan, S., Alessio, N., Acar, M. B., Mber, E., Piber, F., Vitiello, M., & Galderisi, U. (2016). Unbiased analysis of senescence associated secretory phenotype (SASP) to identify common components following different genotoxic stresses. Aging, 8(7), 1316–1329. 10.18632/aging.100971

Roca, H., Hernandez, J., Weidner, S., McEachin, R. C., Fuller, D., Sud, S., Schumann, T., Wilkinson, J. E., Zaslavsky, A., Li, H., Maher, C. A., Daignault-Newton, S., Kuick, R., & Pienta, K. J. (2013). Transcription factors OVOL1 and OVOL2 induce the mesenchymal to epithelial transition in human cancer. PLoS One, 8(10), e76773.

Roychoudhuri, R., Clever, D., Li, P., Wakabayashi, Y., Quinn, K. M., Klebanoff, C. A., Ji, Y., Sukumar, M., Eil, R. L., Yu, Z., Spolski, R., Palmer, D. C., Pan, J. H., Patel, S. J., Macallan, D. C., Fabozzi, G., Shih, H. Y., Kanno, Y., Muto, A., Zhu, J., Gattinoni, L., O’Shea, J. J., Okkenhaug, K., Igarashi, K., Leonard, W. J., & Restifo, N. P. (2016). BACH2 regulates CD8+ T cell differentiation by controlling access of AP-1 factors to enhancers. Nature Immunology, 17(7), 851–860.

Wang, M., Lei, M., Chang, L., Xing, Y., Guo, Y., Pourzand, C., Bartsch, J. W., Chen, J., Luo, J., WidyaKarisma, V., Nisar, M. F., Lei, X., & Zhong, J. L. (2021). Bach2 regulates autophagy to modulate UVA-induced photoaging in skin fibroblasts. Free Radical Biology and Medicine, 169, 304–316. 10.1016/j.freeradbiomed.2021.04.003

Zou, Z., Long, X., Zhao, Q., Zheng, Y., Song, M., Ma, S., Jing, Y., Wang, S., He, Y., Esteban, C. R., Yu, N., Huang, J., Chan, P., Chen, T., Izpisua Belmonte, J. C., Zhang, W., Qu, J., & Liu, G. H. (2021). A single-cell transcriptomic atlas of human skin aging. Developmental Cell, 56(3), 383–397.

Huang, S., Eichler, G., Bar-Yam, Y., & Ingber, D. E. (2005). Cell fates as high-dimensional attractor states of a complex gene regulatory network. Physical Review Letters, 94(12), 128701.

Huang, S., Ernberg, I., & Kauffman, S. (2009). Cancer attractors: a systems view of tumors from a gene network dynamics and developmental perspective. Seminars in Cell & Developmental Biology, 20(7), 869–876.

Kauffman, S. A. (1969). Metabolic stability and epigenesis in randomly constructed genetic nets. Journal of Theoretical Biology, 22(3), 437–467.

Kauffman, S. A. (1993). The Origins of Order: Self-Organization and Selection in Evolution. Oxford University Press.

Kimmel, J. C., Penland, L., Rubinstein, N. D., Hendrickson, D. G., Kelley, D. R., & Rosenthal, A. Z. (2019). Murine single-cell RNA-seq reveals cell-identity- and tissue-specific trajectories of aging. Genome Research, 29(12), 2088–2103. 10.1101/gr.253880.119

Li, C., & Wang, J. (2014). Quantifying the underlying landscape and paths of cancer. Journal of the Royal Society Interface, 11(100), 20140774.

Lopez-Otin, C., Blasco, M. A., Partridge, L., Serrano, M., & Kroemer, G. (2013). The hallmarks of aging. Cell, 153(6), 1194–1217.

Tabula Muris Consortium. (2020). A single-cell transcriptomic atlas characterizes ageing tissues in the mouse. Nature, 583(7817), 590–595.

Trapnell, C., Cacchiarelli, D., Grimsby, J., Pokharel, P., Li, S., Morse, M., Lennon, N. J., Livak, K. J., Mikkelsen, T. S., & Rinn, J. L. (2014). The dynamics and regulators of cell fate decisions are revealed by pseudotemporal ordering of single cells. Nature Biotechnology, 32(4), 381–386. 10.1038/nbt.2859

